# Microbial community assembly during aging of food waste-derived hydrochar: key bacterial guilds mediate nutrient dynamics

**DOI:** 10.1101/2024.02.16.580608

**Authors:** Yang Ruan, Ziyan Wang, Shiyong Tan, Hao Xu, Liyue Wang, Lixuan Ren, Zhipeng Liu, Shiwei Guo, Qirong Shen, Guohua Xu, Ning Ling

## Abstract

Hydrochar aging may mitigate the adverse effects of direct application on plant growth. However, little is known about the strategies for accelerating this process and the microbiological mechanisms involved. This study explored the effects of three strategies, including the additions of straw and efficient-degrading microorganisms, on the aging processes of food waste-derived hydrochar, and the key microbial drivers. The bacterial and fungal community compositions in hydrochar were estimated at different aging periods (i.e., day 1, 7, 14 and 35) using 16S and ITS rRNA gene sequencing. The results showed that straw and microbial inoculum addition improved the temperature and shortened the aging time by ∼30%. Three bacterial guilds, mainly including *Bacillus*-like species, were identified that had significant correlations with the nitrogen (N) and phosphorus (P) dynamics during aging. This study demonstrates the feasibility of manipulating key microbial guilds artificially to achieve efficient harmless-treatment of hydrochar.

**Highlights:**

- Straw and microbial inoculum addition reduced ageing duration by 30%.
- Microbial inoculum addition increases the aging reaction temperature by 13%.
- Bacteria, rather than fungi, play the critical role in hydrochar aging.
- *Bacillus* and related species drive nitrogen and phosphorus cycles of aging.

**Graphical Abstract:** 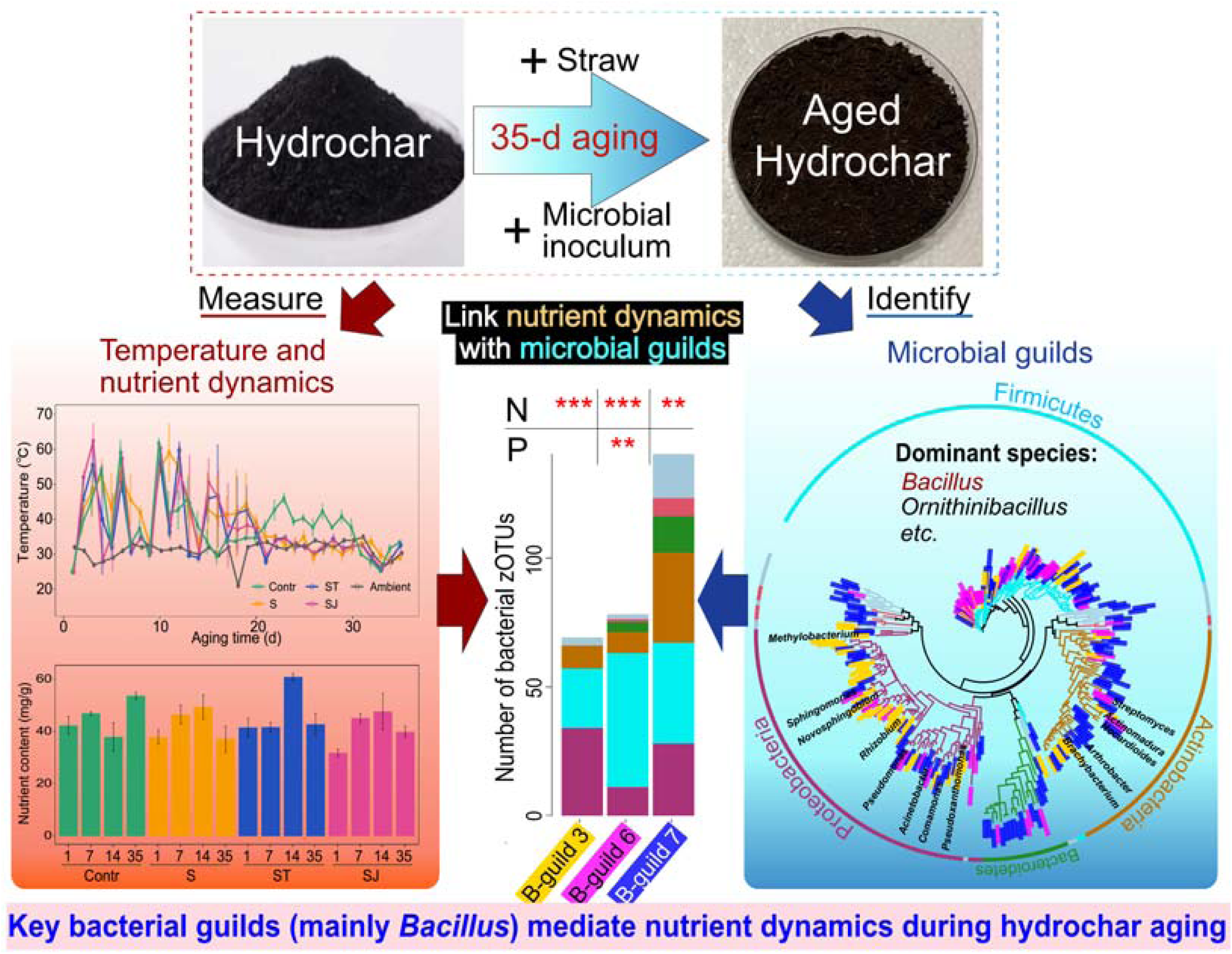

## 1. Introduction

Recovering materials and energy from waste biomass has drawn widespread attention due to the growing population and demand for energy and resources in the past few decades (Dutta et al., 2021; Esquivias et al., 2023). Food waste generated during daily food processing or consumption accounts for more than 50% of all urban organic waste (Zhang et al., 2021a; Grinberga-Zalite and Zvirbule, 2022), which is becoming a critical environmental issue (Forbes et al., 2021). Hydrothermal carbonization has attracted considerable attention as a cleaner production process (Yu et al., 2019). Compared with conventional disposal approaches for food waste, e.g., incineration and landfilling, hydrothermal treatment can convert moisture-rich biomass to hydrochar or biocrude at 180-260 L in a few hours without the need for pretreatment (Zhang et al., 2021b), reducing the cost of hydration (Shen et al., 2021; Yu et al., 2019). The generated lignite-like hydrochar can be used as a conditioner for the improvement of soil structure, fertility, and microbial activity.

Direct application of newly produced hydrochar may have negative effects on plants, including reduced plant biomass, pollen aberration and inhibition of seed germination (Busch et al., 2013; Schimmelpfennig et al., 2014). This is mainly due to 1) the production of derived phytotoxic organic constituents, e.g., furans, polyaromatic hydrocarbons and carboxylic acids (Busch et al., 2012; Karatas et al., 2022), and 2) nitrogen immobilization caused by the high C:N ratio of hydrochar decreasing plant N availability (Bargmann et al., 2014; Gajić and Koch, 2012). Aging process can alleviate the phytotoxic behaviour of hydrochar (Karatas et al., 2022; Yu et al., 2020). For example, cocompost of hydrochar and microbially active compost eliminated the negative effects on cabbage seed germination and plant growth, with similar effects on French marigolds (Roehrdanz et al., 2019). This result suggested that the organic compounds responsible for the negative effects are biodegradable (Busch et al., 2013). However, the aging process often takes tens (Busch et al., 2013) or even hundreds of days (Guo et al., 2014; Wang et al., 2021), which severely limits the application of hydrochar in production. Strategies to accelerate the ageing process and material cycling are therefore urgently needed.

Microorganisms, as decomposers, play the critical role in decomposition of organic matter and release of plant-available nutrients (Karatas et al., 2022). Previous studies demonstrated that the application of microbially aged hydrochar can provide macroelement nutrients for plant growth, increasing the grain yield and free amino acid content (Karatas et al., 2022; Yu et al., 2019). Bacteria may contribute to the nutrient dynamics of hydrochar during aging. In the initial phase of composting, several mesophiles, such as *Escherichia coli* and *Klebsiella* (belonging to Proteobacteria), broke down organic compounds by exothermic reactions; As the temperature rises above 40 L, these mesophiles will be outcompeted by more thermophilic bacteria such as *Bacillus stearothermophilus* (belonging to Firmicutes); The thermophilic bacteria will continue the decomposition processes and produce plant-available nutrients (Insam and de Bertoldi, 2007). Additionally, Fungi generally have a higher carbon use efficiency than bacteria (Tian et al., 2024), which can produce large amounts of mycelium and exoenzyme for nutrient acquisition (Tian et al., 2024), and have a high tolerance for extreme conditions (e.g., high temperature and high salt concentrations) (Gostinčar et al., 2009). Large substrate availability was found to favoured fungal growth, leading to higher fungal biomass and fungal/bacterial ratios (Tian et al., 2024). However, bacterial and fungal community assembly and their contribution to nutrient dynamics during hydrochar aging remain largely unknown.

Different aging strategies would affect the microbial properties, and further influence the aging processes and nutrient contents of aged hydrochar. Here, a 35-day aging experiment with three aging strategies, aging with straw addition (S), aging with straw and decomposition agent addition (ST) and aging with straw and microbial consortium addition (SJ), were performed. Hydrochar samples were collected dynamically over time (Day 1, Day 7, Day 14 and Day 35 of aging) for the chemical properties measurement and 16S and ITS rRNA gene sequencing. The aims of this study are to explore 1) the effects of different strategies on the aging processes, including temperature and duration of reaction; 2) the responses of bacterial and fungal community assembly to different aging strategies and 3) potential key microbial guilds contributing nutrient dynamics during aging. The results can lay the foundation for improvement of aging efficiency and product quality of food waste-derived hydrochar by microbial inoculum.

## 2. Materials and methods

### 2.1. Preparation of food waste-derived hydrochar

Food waste is collected from the cafeteria of Nanjing Agricultural University, which generates about 150 tons of food waste per month. To minimize the variation of sampling, food waste samples were collected in four quarters of the year and stored at −4 L. Hydrochar was prepared from food wastes in a 50 L self-pressured hydrothermal reactor with a 9 kW electrical heater manufactured by Chaoyang Chemical Engineering Company (Shandong, China). 15 kg food waste was added to the reactor for each reaction. The reaction was carried out at a temperature of 180 L, and the retention time was kept at 60 min. The reactor was cooled naturally at the end of the reaction. Then the mixtures flowed out through a valve, and solid-liquid separation was carried out using a filter press with gauze. The hydrochar was dried at 105L for 24 h and subsequently subjected to an aging experiment.

### 2.2. Preparation of microbial consortium

The strain was expanded before the aging experiment. The microbial consortium is composed of four strains of *Bacillus subtilis*, *Aspergillus fumigatus*, *Thermomyces lanuginosus*, and *Trichoderma guizhouense* mixed in a ratio of 1:1:1:1. All these strains were the dominant strains during straw composting and were isolated from the laboratory. *B. subtilis* has several advantages for heterologous protein production, involving high secretion capacity and easy fermentation culture (Zhao et al., 2019). *Aspergillus fumigatus* (Liu et al., 2013), *Thermomyces lanuginosus* (Zhang et al., 2019) and *Trichoderma guizhouense* (Meng et al., 2020) were found to have the strong ability to decompose lignocellulose biomass. All these strains can adapt to high temperature conditions. Toothpicks were used to pick a colony and placed in LB liquid culture medium (PDA medium for fungus). The culture flasks were placed in a shaking table at 28 L and a speed of 200 r/min for 2-3 days (3-4 days for fungus) in the dark. The plate-counting method was used to estimate the total number of bacterial cells or fungal spores (Gorsuch et al., 2019), and the final concentration was higher than 1 × 10^9^ cfu/mL.

### 2.3. Hydrochar aging experiment

The aging experiment was set up with four treatments: normal hydrochar aging with as the main material using water as the solvent (500 g hydrochar + 100 g sterile water, Contr); hydrochar aging with straw addition (500 g hydrochar + 50 g maize straw + 100 g sterile water, S); hydrochar aging with straw and decomposition agent addition (500 g hydrochar + 50 g maize straw + 100 g sterile water + 1% of the decomposition agent, ST); and hydrochar aging with a microbial consortium and straw addition (500 g hydrochar + 50 g maize straw + 100 g sterile water + 1% of the microbial consortium, SJ). The decomposition agent was purchased from a local fertilizer factory.

The aging processes were performed in a 12 L fermenter. The hydrochar was built into piles and flipped every three days. The temperature of each reactor was monitored daily. The aging process was completed when the temperature of the pile did not change for five consecutive days. Hydrochar samples were collected on Days 1, 7, 14 and 35 (the end of aging). 15 g hydrochar was sampled from each sampling and divided into two parts: one was stored at 4 L for total microbial DNA extraction, and the other was air dried to measure the chemical properties.

### 2.4. Measurement of chemical properties

The measurement methods were similar to those reported in a previous study (Guo et al., 2020). Briefly, the pH and Ec of hydrochar were determined in hydrochar:water (1:2.5, w/v) by a combination electrode (PE-10, Sartorious, Germany). The nutrients (i.e., nitrogen, phosphorus, and potassium) from hydrochar were released by using the H_2_SO_4_-H_2_O_2_ digestion method (Guo et al., 2021). The nitrogen and phosphorus contents were determined by an AutoAnalyzer (AA3, Bran+Luebbe, Germany). The potassium contents were determined by flame photometry (AP1200; Shanghai Aopu Analytical Instruments, Shanghai, China).

### 2.5. DNA extraction, bacterial 16S rDNA, and fungal ITS2 sequencing

DNA was extracted from 500 mg of hydrochar sample using the FastDNA™ SPIN Kit for Soil (MP Biomedicals, Cleveland, OH, USA) according to the manufacturer’s instructions. The V3-V4 regions of the bacterial 16S rRNA genes were amplified using primers 341F (5’-CCTACGGGNGGCWGCAG-3’) and 805R (5’-GACTACHVGGGTATCTAATCC-3’) (Pan et al., 2023). The internal transcribed spacer 2 (ITS2) regions of the fungal ribosomal RNA gene were amplified using primers ITS1FI2 (5’-GTGARTCATCGAATCTTTG-3’) and ITS2 (5’-TCCTCCGCTTATTGATATGC-3’) (Männistö et al., 2018). The amplicons were sequenced with the Illumina MiSeq^TM^ system. The raw sequences were uploaded to the National Genomics Data Center (NGDC) Genome Sequence Archive (GSA) with accession number CRA012057.

The sequences were quality-filtered and analysed using USEARCH v.11.0 (Edgar, 2010). Forwards and reverse reads were merged, and low-quality sequences (i.e., length < 200 bp, quality score < 20) were filtered. After exclusion of chimaeras, the remaining sequences were clustered into zero-radius operational taxonomic units (zOTUs) by the Unoise3 algorithm (Edgar, 2016). Finally, the representative sequence of each zOTU was matched against the rdp_16s (Cole et al., 2014), and UNITE (Kõljalg et al., 2013) databases for the bacterial and fungal taxonomic assignment, respectively.

### 2.6. Diversity analyses

The richness and Shannon index of bacterial and fungal communities were calculated using the “vegan” package in R 4.2.1 (http://www.r-project.org). Permutational multivariate analysis of variance (PERMANOVA) and principal coordinate analysis (PCoA) were used to compare the beta diversity of bacteria and fungi based on the Bray-Curtis distance matrix by the “ape” and “vegan” packages in R 4.2.1(Ruan et al., 2020). The microbial co-occurrence network was constructed to infer the interspecies interactions. Network analysis at the bacterial and fungal zOTU levels was performed using the R package WGCNA (Langfelder and Horvath, 2008). The microbial co-occurrence network of each aging treatment was characterized and visualized using Gephi (v0.9.5) (Ruan et al., 2020). The normalized stochasticity ratio (NST) of community assembly was calculated based on Bray-Curtis metrics using the null model algorithm (R package NST) (Ning et al., 2019). The NST index was developed with 50% as the boundary point between more deterministic (< 50%) and more stochastic (> 50%) assembly (Ning et al., 2019). Differential zOTU analysis (using the R package DESeq2) was performed to identify the significantly enriched/depleted OTUs in the three aging strategies with straw addition compared to the Contr treatment (Ruan et al., 2020). The results of differential zOTU analysis were visualized using heatmap analysis (R package “pheatmap”). Bacterial and fungal guilds were further divided based on the fold change of the differential abundance species (Log_2_FC) in the comparison between each aging strategy and the Contr treatment during the aging periods by using WGCNA (Liu et al., 2020). Linear regression analyses were conducted to estimate the relationship between the abundance of microbial guilds and the nutrient contents (R 4.2.1, ‘lm’ function) (Ruan et al., 2023). The phylogenetic tree of the taxa in the potential key guilds was structured in Galaxy/DengLab (http://mem.rcees.ac.cn:8080/) with PyNAST Alignment and FastTree functions (Caporaso et al., 2009; Price et al., 2009) and visualized by the Interactive Tree Of Life (iTOL) (Letunic and Bork, 2021). KO gene annotation of taxa was performed by PICRUSt2 analysis (Douglas et al., 2020). The indicator genes related to carbon (Llorens-Marès et al., 2015) and nitrogen (Llorens-Marès et al., 2015; Nelson et al., 2015), sulfur (Llorens-Marès et al., 2015), and phosphorus (Dai et al., 2019) cycling were selected according to the conclusions reported in previous publications.

## 3. Results and discussion

### 3.1. Dynamics of temperature and nutrient contents under aging strategies

Temperature changes are closely related to the microbial activity in the reactor, which works to decompose the organic substrates and generate heat energy (Liu et al., 2023). Thus, the changes in temperature can be regarded as an important index for evaluating the reaction state of aging. The results showed that the hydrochar aging process consists of four obvious heating reactions (Fig. 1a). The first thermal reaction occurred within five days of aging: The temperature of the hydrochar samples increased rapidly after one day of aging. The temperature of the hydrochar in the SJ and ST treatments reached the first peak on Day 3 (62 L and 55 L, respectively). After four days of aging, the temperature in Contr and S reached the first peak (54 L and 52 L, respectively), and the temperature in SJ decreased to a minimum (28 L). Then, the temperature of the other three treatments decreased to a minimum on Day 5. At the second thermal reaction, the temperature in all four aging treatments reached the second peak on Day 6 and dropped to a minimum on Day 9. The third thermal reaction of the S, ST and SJ treatments lasted from Day 9 to Day 14, while the Contr treatment lasted from Day 9 to Day 16. Finally, the fourth thermal reaction of the S, ST and SJ treatments lasted from Day 14 to Day 21, while the Contr treatment lasted from Day 20 to Day 31.

**Fig. 1.**
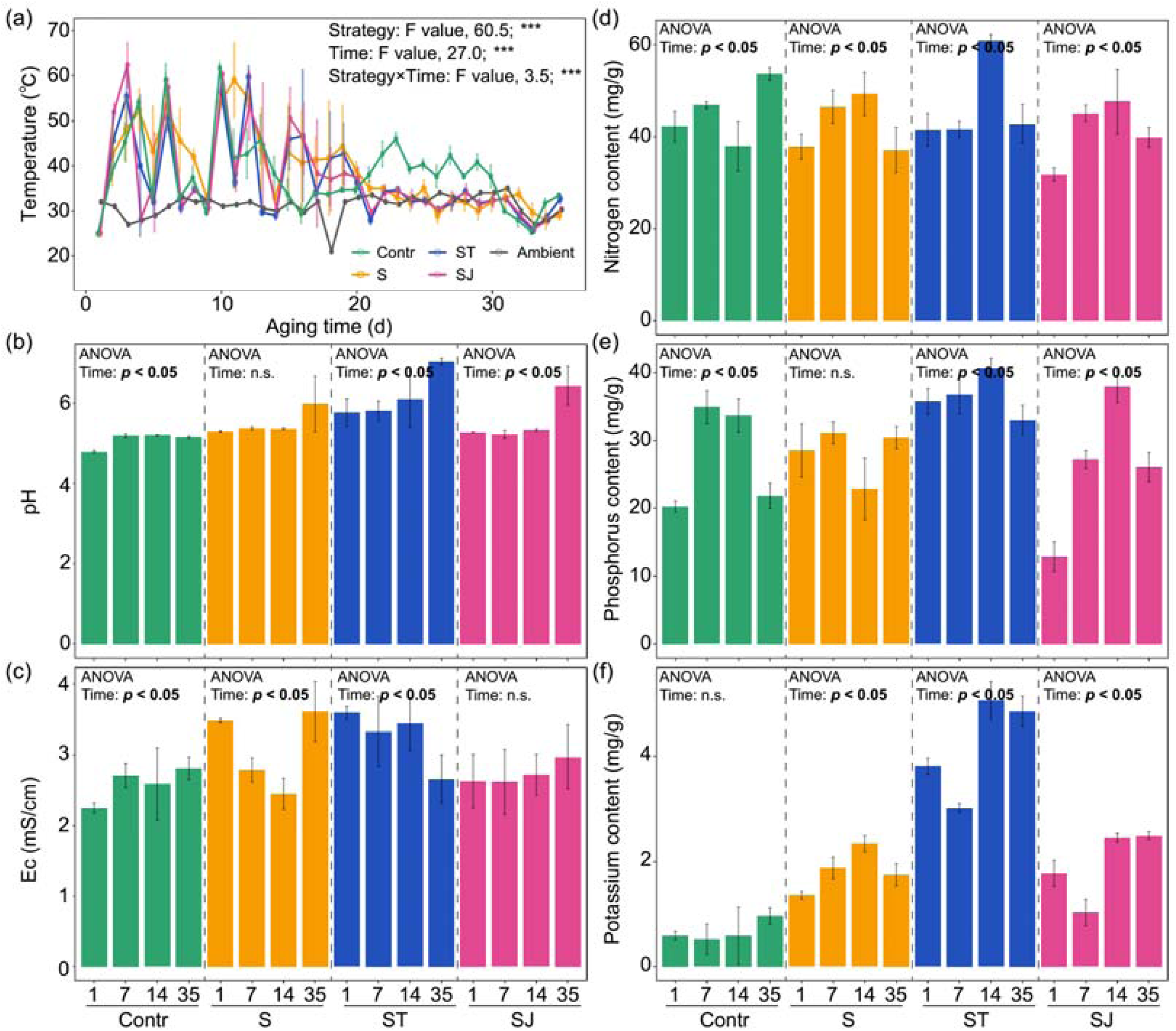
Dynamics of temperature and chemical characteristics during the aging process of hydrochars. The dynamics of temperature during the 35 d aging processes under the aging strategies **(a)**. Changes in pH **(b)**, Ec **(c)**, nitrogen content **(d)**, phosphorus content **(e)**, and potassium content **(f)** under different periods of the four aging treatments. Data are presented as the mean values ± SDs (n = 3). Contr represents the normal aging treatment; S represents the hydrochar aging with straw addition; ST represents the hydrochar aging with straw and decomposition agent addition; and SJ represents the hydrochar aging with straw and the microbial consortium addition. *** represents “*p* < 0.001”; n.s. represents that there is no significant difference among treatments.

In the initial stages of hydrochar aging, there was an increasing trend in the compost temperature for all aging treatments, while the addition of the decomposition agent and microbial consortium made the temperature peak one day earlier and at higher temperatures (approximately 7-17L) than those in the Contr and S treatments (Fig. 1a). Higher temperatures are beneficial for killing harmful microorganisms such as *Ascaridia galli* eggs, *Escherichia coli* and *Salmonella enterica* in the raw hydrochar material (Chen and Jiang, 2017; Katakam et al., 2014). Furthermore, the temperature in the SJ last peaked on the 15th day of aging, four days earlier than that in the S and ST, while the temperature in the Contr treatment was obviously higher than the ambient temperature until the 31st day of aging. This result suggested that the addition of straw and the microbial consortium accelerated the aging process of hydrochar.

Hydrochar samples from different aging stages were collected for basic chemical analysis (Fig. 1b-f). The three aging strategies with straw addition significantly increased the pH of the hydrochar samples (ANOVA, *p* < 0.05). The aging time also affected the pH of the hydrochar: the pH significantly increased with aging time, except for the S treatment (*p* > 0.05, Fig. 1b). The Ec of hydrochar was also affected by aging strategy and aging time. The dynamics of hydrochar nutrient contents (i.e., nitrogen, phosphorus, and potassium concentrations) depended on aging time (Fig. 1d-f). The nitrogen and phosphorus concentrations first increased and then decreased, and the potassium concentration increased with aging time. These results indicate that the chemical composition of hydrochar changed during the aging process.

The pH level played a crucial role in the activity of microorganisms (Rousk et al., 2010). The results showed that the level of pH increased with aging time for all treatments (Fig. 1b), which could be due to NH_4_^+^-N generated through the mineralization of organic matter (Li et al., 2023). Ec denotes the level of compost salinity and maturity. The Ec increase could be caused by the release of mineral salts such as phosphates and ammonium ions through the decomposition of organic substances, while the volatilization of ammonia and the precipitation of mineral salts could decrease the level of Ec (Gao et al., 2010; Sun et al., 2021). Ec values over 4 mS/cm are usually detrimental to plant growth (Li et al., 2023). In this study, the Ec values of hydrochar, in all aging treatments, were below 4 mS/cm, indicating limited toxic effects on plants.

At the end of aging, the nitrogen and phosphorus contents of hydrochar did not decrease significantly compared with the samples at the beginning of aging, indicating limited nutrient losses (Fig. 1d and e). Microorganisms release large amounts of nitrogen by organic mineralization, and the excess N can be lost from the hydrochar through leaching or volatilization as ammonia (Gao et al., 2010). Moreover, water evaporation owing to the heat generated through microbial activity was the primary cause of moisture content loss (Li et al., 2023), causing an increase in nutrient content per unit of hydrochar during the aging process. This finding could explain the observed increase in the potassium concentration of hydrochar during the aging periods (Fig. 1f).

### 3.2. Diversity of bacterial and fungal communities under aging treatments

The richness and Shannon indices reflected the magnitude of species number and diversity, i.e., alpha diversity. The results showed that the bacterial alpha diversity (both richness and Shannon indices) of the hydrochar samples was significantly higher than the fungal diversity at each aging stages (Table S1). The dramatic difference in alpha diversity between bacterial and fungal communities indicated that the aging process is highly selective and favourable to bacterial species, which could be attributed to the faster reproductive rate and shorter reproductive time of bacteria (Tian et al., 2024). The aging time had slight effects on bacterial and fungal richness, with the exception of the bacterial community in the SJ (significantly decreased with aging time). The dynamics of Shannon indices during aging remained stable in the bacterial communities. These results suggested that hydrochar aging time has a limited impact on microbial alpha diversity.

The variations in species compositions (i.e., beta diversity) of the hydrochar bacterial and fungal communities were further estimated among aging treatments at each aging period (Fig. 2 and 3). The phylum-level compositions of bacterial communities at the initial stage of aging were similar among the aging treatments (Fig. 2a), mainly composed of Proteobacteria (53%) and Firmicutes (26%). Similar to this result, previous studies found that the microbial community in compost was mainly composed of Proteobacteria, Firmicutes, and Actinobacteria (Bao et al., 2023; Li et al., 2023). Proteobacteria contain numerous metabolic types and constitute one of the most diverse and abundant groups of microbes (Zhou et al., 2020). Members of Firmicutes play important roles in plant growth promotion and pathogen biocontrol (Gavande et al., 2021; Zhu et al., 2016). The relative abundance of bacterial phyla varies noticeably during the aging process. After 7 days of aging, the abundance of the Proteobacteria phylum decreased by 42% in the Contr and ST treatments and increased for Firmicutes in all four aging treatments (85%, 157%, 87% and 8% in Contr, S, ST and SJ, respectively). On Day 14, the abundances of Actinobacteria and Bacteroidetes increased in the Contr (6.6 and 3.5 times, respectively) and ST (3.6 and 1.8 times, respectively). The abundance of Firmicutes decreased by ∼50% in the Contr and ST and increased by ∼75% in the S and SJ. At the end of aging, Proteobacteria dominated the bacterial communities of hydrochar in the Contr and ST treatments (with relative abundances of 54% and 83%, respectively), while Firmicutes became dominant in the S treatment (61%). The species composition at the phylum level was more uniform in the SJ treatment than in the other three treatments at the end of aging: The abundance of Firmicutes was the highest (36%), followed by that of Proteobacteria (34%), Actinobacteria (15%) and Bacteroidetes (13%).

**Fig. 2.**
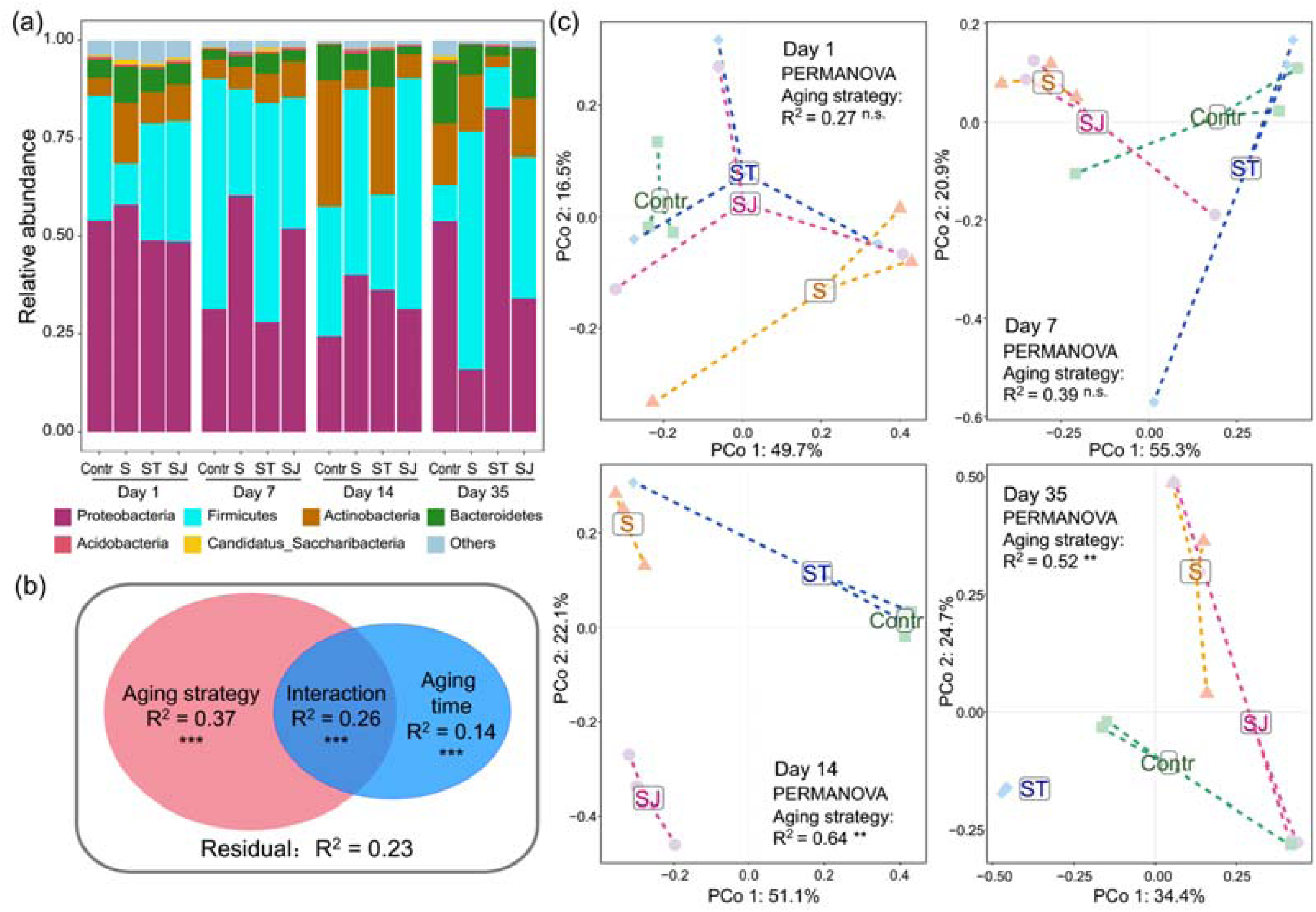
Hydrochar bacterial communities are separable by aging strategy and aging time. Taxonomic composition of the bacterial community at the phylum level **(a)**. Two-way permutational multivariate analysis of variance (PERMANOVA, **b**) and principal coordinate analysis (PCoA) plot **(c)** depict the Bray–Curtis distance of the bacterial community.

**Fig. 3.**
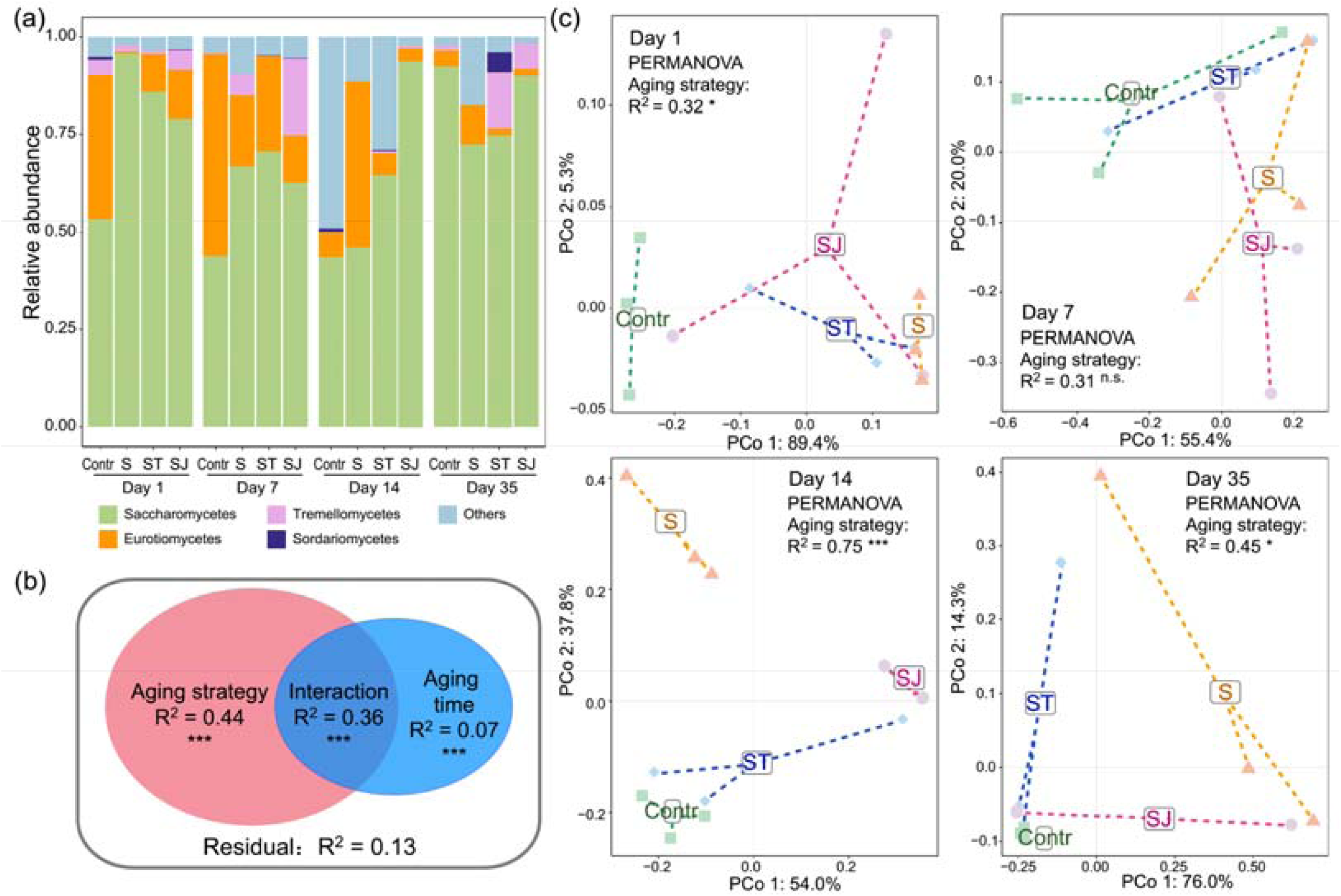
The dynamics of hydrochar fungal communities under the aging strategies. Taxonomic composition of the fungal communities at the phylum level **(a)**. Two-way PERMANOVA **(b)** and PCoA plot **(c)** depict the Bray–Curtis distance of the fungal communities.

Both aging strategy and aging time significantly influenced the bacterial compositions of hydrochar (PERMANOVA, both *p* < 0.001), where the effect of aging strategy (PERMANOVA, R^2^ = 0.37) was greater than that of aging time (PERMANOVA, R^2^ = 0.14, Fig. 2b). Principal coordinate analysis (PCoA) were performed to visualize the difference in bacterial species composition between aging treatments (Fig. 2c). There was no significant difference in species composition among aging treatments after 1-week aging (*p* > 0.05). After a 2-week aging period, the bacterial composition significantly diverged due to the aging strategy (R^2^= 0.64, *p* < 0.01), which persisted until the end of the aging process (R^2^= 0.52, *p* < 0.01). These results suggested that aging strategy and aging time have a slight effect on alpha diversity, but a strong effect on beta diversity. Similar to the results of this study, previous research showed that some experimental manipulations (e.g. nutrient addition) may not significantly affect the species diversity (i.e., alpha diversity) of microbial community, but species composition (i.e., beta-diversity). These results suggest that beta-diversity may more sensitively reflect the response of microorganisms to environmental changes. Such changes in microbial communities indicated the different environmental conditions and substrate compositions at different stages of aging (Zhao et al., 2018).

The phylum-level compositions of fungal communities were mainly composed of Saccharomycetes and Eurotiomycetes (average abundances: 71% and 15%, respectively, Fig. 3a). The three aging strategies, on average, increased the abundance of Saccharomycetes by 63% on Day 1 and by 52% on Day 7, and, on average, decreased the abundance of Eurotiomycetes by 80% and by 65%. At the end of aging, Saccharomycetes dominated the fungal communities of hydrochar, while the abundance of Tremellomycetes showed a noticeable increase in the ST and SJ treatments (relative abundance: 14% and 6%, respectively). Similar to the results for bacteria, aging strategy (R^2^ = 0.44, *p* < 0.001) also contributed to the largest variation in fungal communities, followed by aging time (R^2^ = 0.07, *p* < 0.001). The interaction between aging strategy and time also affected the beta diversity of the fungal communities (R^2^ = 0.36, *p* < 0.001, Fig. 3b). The PCoA results showed significant divergence among aging treatments at different periods (except for sampling on Day 7, Fig. 3c). These results suggested that the compositions of the hydrochar microbiome exhibited distinct responses to aging strategies and aging time, which could influence microbial functional performance (Hua et al., 2020; Liu et al., 2023). Therefore, exploring the community assembly of hydrochar microbes would be conducive to elucidating microbe-driven nutrient turnover processes during aging.

### 3.3. Co-occurrence network of the hydrochar microbiome

Co-occurrence network analysis was performed based on the abundance matrix of bacterial and fungal communities to characterize the interactions between microbial taxa, (Fig. 4). The results showed that bacterial species dominated the microbial interaction network in all aging treatments (> 96% network nodes). Similar to the result of alpha diversity, bacteria dominated the interaction networks in all four aging treatments, indicating a potential key role in hydrochar aging. The number of network nodes showed a slight decrease (by 6%), and the edge number decreased dramatically (by 57%) after straw addition (S treatment). The reduction effects on edge number were almost fully alleviated with the addition of decomposed liquids (ST) and microbial consortium (SJ). Bioavailable carbon sources are critical to microbial maintenance and growth (Ruan et al., 2023). Crop straw, as a renewable resource, is extensively used in modern agriculture worldwide, affecting soil nutrient properties (Tang et al., 2020) and microbial communities simultaneously (Ma et al., 2019). Straw contains a large amount of complex carbon sources, such as cellulose and hemicellulose, which are difficult for most microorganisms to utilize. Straw addition altered the material composition of aqueous charcoal, making it less likely that some microbial species that feed on labile carbon will be able to survive, thus reducing network complexity (Hernandez et al., 2021). The level of network complexity was restored to the original state by the addition of the decomposition agent and microbial consortium (Fig. 4). Both of these additions can efficiently decompose complex carbon sources in straw into labile carbon for utilization by other microorganisms, thereby enhancing network interactions.

**Fig. 4.**
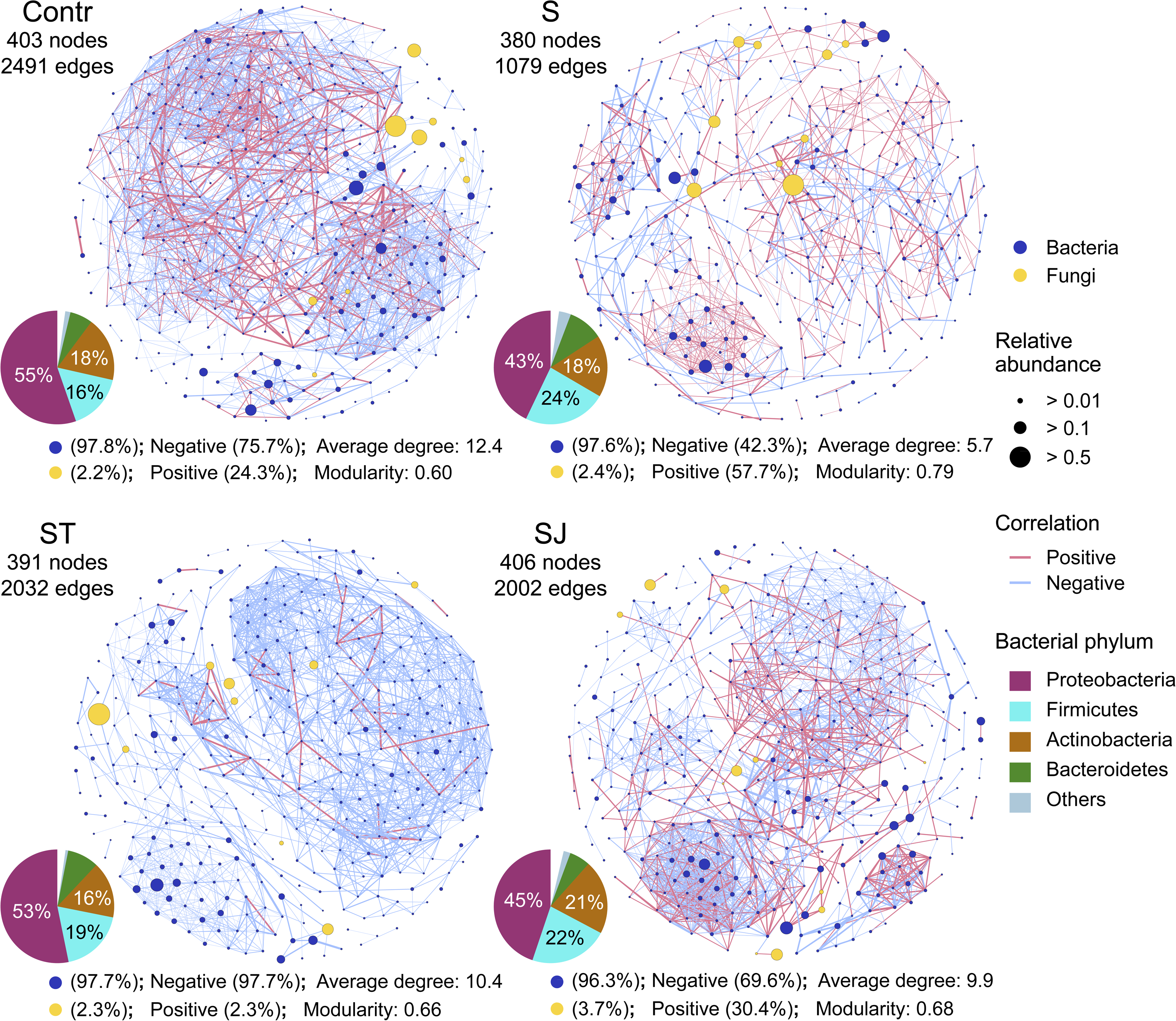
Bacterial-fungal co-occurrence network analysis of hydrochars under the aging strategies. The blue nodes represent bacterial taxa, and yellow nodes represent fungal taxa. The node size indicates the relative abundance. Positive and negative correlations between taxa are shown by different edge colours. The pie chart represents the proportion of nodes belonging to different bacterial phyla in each network.

As the dominance of bacteria in the network, the taxonomic composition of bacterial nodes was further investigated at the phylum level. The nodes of the microbial network in hydrochar consisted mainly of Proteobacteria, Firmicutes, Actinobacteria, and Bacteroidetes. Among the four aging treatment networks, the Contr network had the highest percentage of nodes belonging to Proteobacteria in the network (55%), followed by ST (53%), SJ (45%), and S (43%). Compared to the Contr treatment, the three aging strategies with straw addition increased the proportion of nodes belonging to Firmicutes by 1.2-1.5 times. Changes in microbial network structure affect ecological functions (Yuan et al., 2021; Zamkovaya et al., 2021). The increased proportion of Firmicutes indicated a potential critical role in the three aging processes. Firmicutes was found to be the dominant phylum in replicate nutrient-cycling model ecosystems (Pagaling et al., 2017). Furthermore, members of the Firmicutes, such as *Clostridium thermocellum*, can degrade cellulose by fermenting the mono- and di-saccharide products of cellulose breakdown (Pagaling et al., 2017). It can therefore be concluded that increased proportions of the Firmicutes phylum in the network favoured accelerated substrate decomposition during aqueous charcoal ageing, possibly due to the addition of straw in the three ageing strategies.

### 3.4. Community assembly of the hydrochar microbiome

Community assembly is driven by both deterministic and stochastic processes (Feng et al., 2018; Stegen et al., 2013). The normalized stochasticity ratio (NST) was used to estimate the relative contribution of deterministic and stochastic processes to bacterial and fungal community assembly (Fig. S1). Using 50% as the threshold, an NST greater than 50% means that the stochastic process dominates, and conversely, the deterministic process dominates community assembly (Ning et al., 2019). For the bacterial community, NST was the highest in Contr (91%), while the three aging strategies with straw addition decreased the NST by 61-73% in the initial stage of aging (Fig. S1a). Then, NST rapidly decreased to 34% on Day 7 and dropped to 17% at the end of aging. The dynamics of NST in the S and SJ treatments exhibited an inverted U-shape: NST reached a maximum on Day 7 in S (89%) and peaked on Day 14 in SJ (68%). The NST of all aging treatments was less than 50% at the end of aging, with S (44%), ST (27%), SJ (22%), and Contr (17%) in descending order.

For the fungal communities, NST in Contr and SJ maintained a high level in the first week of aging (70-78%) and then sharply decreased to 11-15% until the end of aging (Fig. S1b). NST in S and ST was less than 50% on Day 1 (5% and 31%, respectively), peaked on Days 7 and 14 (61-74%), and then dropped to below 50% at the end of aging (39% and 29%, respectively).

Disentangling the relative importance of determinism and stochasticity in community assembly is one of the greatest challenges in microbial ecology (Ning et al., 2019; Ruan et al., 2022). Deterministic processes generally refer to any ecological process that involves nonrandom, niche-based mechanisms, including environmental filtering (e.g., pH, moisture, salinity, and nutrient level) and biotic interactions. Stochastic processes typically include random birth-death events, species migration (e.g., random chance for colonization), and ecological drift (Nemergut et al., 2013; Stegen et al., 2013). Previous studies suggested that nutrient level improvement (e.g., organic fertilization) usually increases the proportion of stochastic processes because of the mitigation of nutrient limitations (Feng et al., 2018; Ruan et al., 2023). Compared with the Contr treatment, the three aging strategies dramatically increased the proportion of deterministic processes of the bacterial community at the initial stage of aging. This result indicated that straw addition could cause localized nutrient limitations on bacterial communities because of the large number of recalcitrant and complex carbon sources in straw. The addition of decomposition agents and microbial consortium could release nutrients by decomposing straw, which partially alleviated nutrient limitation, as evidenced by smaller deterministic proportions than that in the S treatment.

As aging progressed, the proportion of stochastic processes increased across treatments (except for the bacterial community in the ST treatment), which may be due to nutrient release by microbial-mediated mineralization of organic matter. The proportion of stochastic processes decreased to less than 50% by the end of aging, probably due to the increased role of environmental selection as a result of reduced microbially available resources during decomposition. Additionally, water evaporation due to the high temperatures during aging would result in limitations on diffusion, which hindered the intimate contact between the organism and substrate. Limited substrate diffusion reduced substrate bioavailability, as insoluble and adsorbed substrates cannot cross the cytoplasmic membrane (Barnett et al., 2021). This can also lead to microbial community assembly that is more strongly influenced by deterministic processes.

Interestingly, at the end of aging, the bacterial community in the S treatment had the highest proportion of stoichiometric processes among the four treatments, which may be because some undecomposed organic matter remained available for microbial utilization. The result of the stochastic ratio in the fungal communities was similar among aging treatments, i.e., rising then falling. Such dynamics of stochastic ratio could be explained by the degradation of complex substrates and changes in nutrient content during aging. Similarly, previous findings indicated important connections among initial conditions, degree of change in environmental variables and microbial community assembly processes, and stochastic processes increase with higher nutrient condition (Feng et al., 2018). Collectively, these results indicated that the degradation of organic matter could be an important driving factor of microbial community assembly.

### 3.5. Enriched and depleted microbial species in the three aging strategies

The processes of organic matter degradation and nutrient transformation are often accompanied by changes in the abundance of associated functional microbial taxa (Hartman et al., 2017). To identify the species with significant variations in abundance by aging strategy, the fold change in species abundance was calculated by comparing the abundance of each zOTU in the three aging strategies with that in the C treatment during the aging periods (Fig. 5). A total of 1219 bacterial species (mainly belonging to Proteobacteria, Firmicutes, Actinobacteria, and Bacteroidetes) and 62 fungal species (mainly belonging to Eurotiomycetes, Sordariomycetes, and Saccharomycetes) were found with differential abundance. The obvious enrichment effects occurred by three aging strategies at the initial stage of aging (Fig. 5a), i.e., more enriched zOTUs than depleted zOTUs in each comparison (1.7-3.6 times), indicating species activation. The enrichment effects persisted in S and SJ until 14 days of aging. At the end of aging (35 d), the number of depleted species was consistently greater than that of enriched species (3.3, 1.5, and 8.0 times in S, ST and SJ, respectively). The depletion effects indicated that the abundance of most bacterial taxa decreased at the end of aging because of the reduction in bioavailable C sources.

**Fig. 5.**
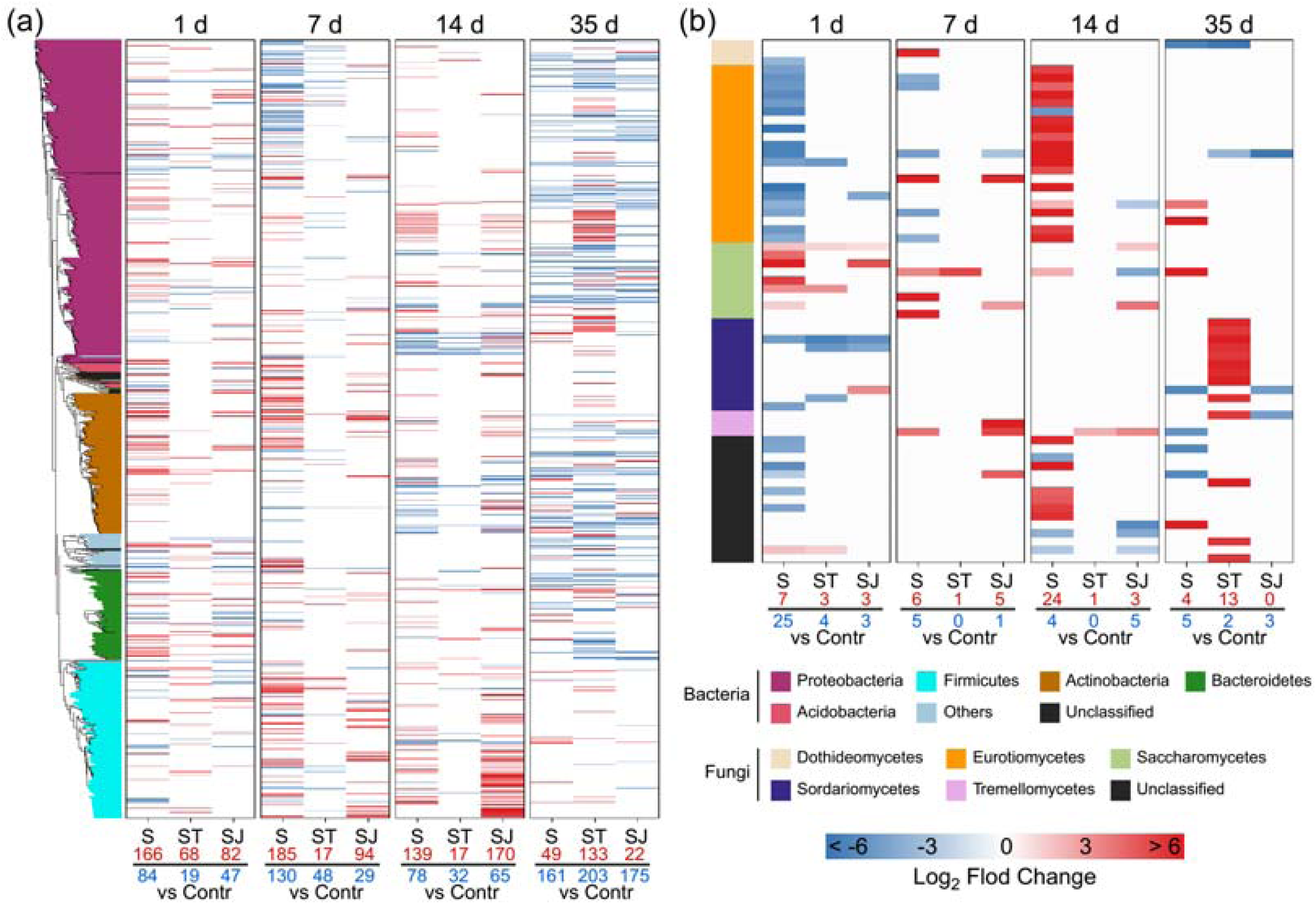
The differential abundance taxa of bacteria (a) and fungi (b) between the three aging strategies (S, ST, and SJ) and the normal aging treatment (Contr) during the aging periods. The heatmap shows the log_2_ fold change of taxa under each comparison. The red number represents the number of enriched zOTUs in the corresponding aging strategy, while the blue number represents the number of enriched zOTUs in the Contr treatment.

The number of fungal species with significant changes in abundance is much smaller than that of bacterial species. Aging with only straw addition (S) mainly depleted the species affiliated with Eurotiomycetes (16 species) and enriched six species affiliated with Saccharomycetes, while the decomposition agent and microbial consortium addition erased this effect (Fig. 5b). Aging with only straw addition (S) enriched 16 species affiliated with Eurotiomycetes after 14 d of aging, and ST enriched 10 species affiliated with Sordariomycetes at the end of aging (35 d).

### 3.6. Key bacterial guilds explains the dynamics of nutrient contents

It is reasonable to assume that these differential abundance species within the three strategies possess important functions such as organic matter decomposition and nutrient transformation. The taxa with differential abundance were widely distributed on phylogenetic trees (Fig. 5), indicating that phylogenetic relatedness was a weak and inconsistent predictor of the microbial abundance dynamics in hydrochar. An alternative means was used to group microbial taxa into ecologically relevant clusters based on the characteristics of species abundance changes (i.e., log_2_ fold change). The differentially abundant species were grouped into 9 bacterial guilds and 2 fungal guilds by WGCNA (Fig. S2). Each bacterial guild contained between 64 and 230 species, while two fungal guilds contained 23 and 39 species. The abundance dynamics of guilds were further related to the variations in nutrient contents during hydrochar aging. The results of the regression analysis showed significant positive correlations between three bacterial guilds and the N and P nutrient dynamics of hydrochar, while no significant correlations were found between fungal guilds and nutrient dynamics (Fig. 6a). Bacterial guild 3, guild 6 and guild 7 were correlated with the nitrogen content, and bacterial guild 6 was also correlated with the phosphorus content of hydrochar (*p* < 0.01). No bacterial or fungal guilds showed significant correlations with the potassium content during aging (*p* > 0.05).

**Fig. 6.**
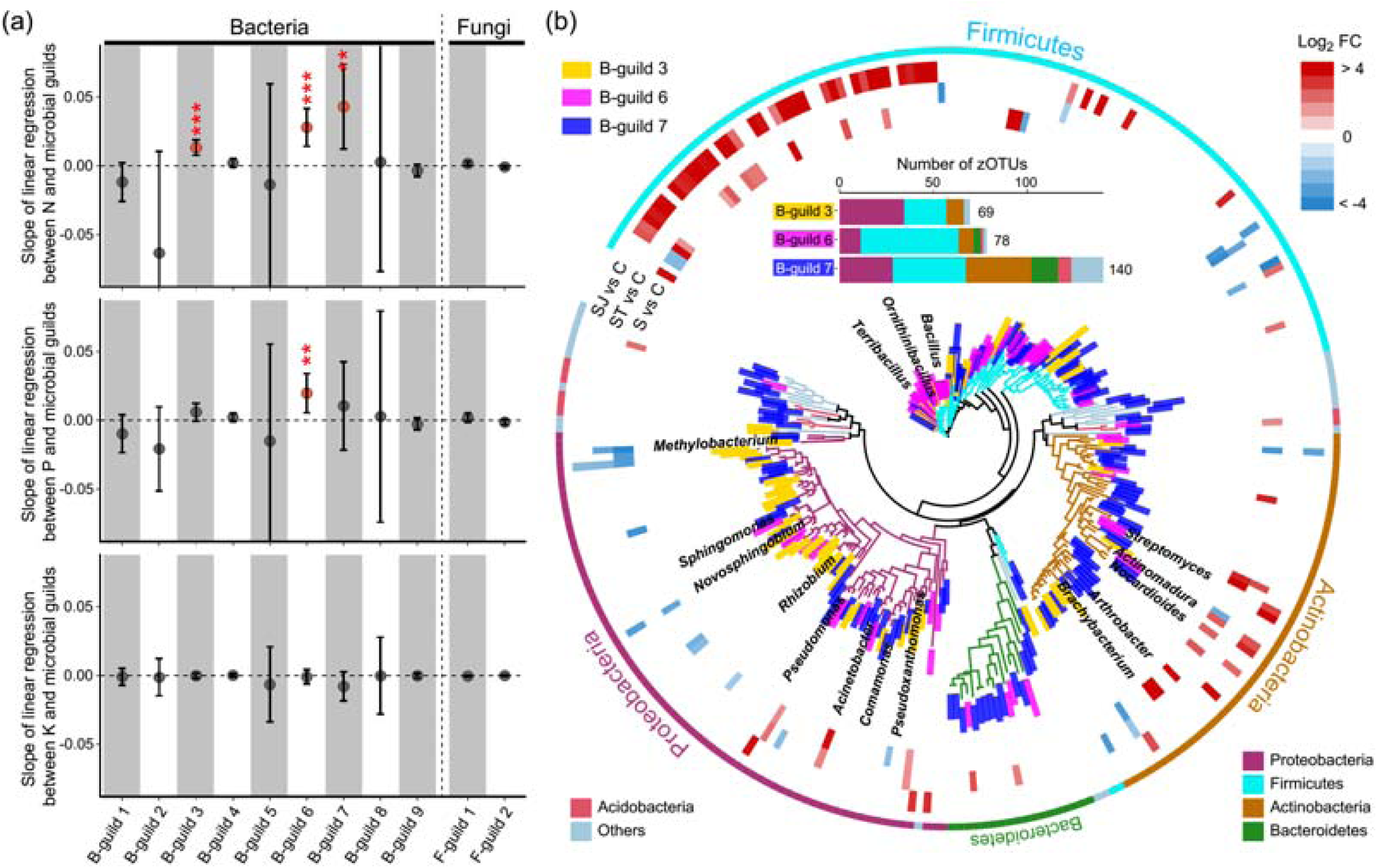
Relevance of microbial guilds to nutrient dynamics during aging processes and the phylogeny of the species from the three key bacterial guilds. Slopes of linear regressions between the nutrient contents (nitrogen, phosphorus, and potassium) of hydrochar and the abundance of microbial guilds during the 35-d aging periods **(a)**. The dots indicate the means of the slopes, and the bars show the 95% confidence intervals. The red asterisk represents a significant regression. The phylogeny of the three bacterial guilds that had significant regression relationships with the nutrient contents of hydrochar **(b)**. The heatmap in the outer circle shows the log_2_ fold change of taxa under each comparison (from the inside out: S vs. Contr, ST vs. Contr, SJ vs. Contr). The stacked diagram represents the number of species in the guild and their classification at the phylum level.

A phylogenetic tree including all three bacterial guilds correlated with nutrient dynamics was constructed, and the species composition of each guild was analysed and visualized (Fig. 6b). Among the three bacterial guilds, guild 3 harboured the most species affiliated with Proteobacteria (34 species), guild 6 harboured the most species belonging to Firmicutes (52 species), and guild 7 had the most species belonging to Actinobacteria (35 species) and Bacteroidetes (14 species). The species composition in guild 7 was more even than that in guilds 3 and 6, mainly composed of 28% Firmicutes, 25% Actinobacteria, 20% Proteobacteria and 10% Bacteroidetes. Conversely, guild 6 was dominated by Firmicutes (proportion of species composition: 67%). Members of Firmicutes are globally distributed and are known to play an important role in the cycling of C, N, P, and S elements (Delgado-Baquerizo et al., 2017; Wasmund et al., 2017). Furthermore, at thermophilic temperatures (approximately 50-60 L), members of Firmicutes are reported to be the major contributors to the conversion of organic matter (Bao et al., 2023; Wrighton et al., 2008), which could be due to metabolic adaptations and the formation of wet-heat resistant spores (Filippidou et al., 2016). As bacterial guild 6 was positively correlated with the dynamics of N and P nutrients (Fig. 6a), it can be inferred that members of Firmicutes are important participants in nutrient cycling during hydrochar aging.

The abundance fold change of each zOTU in the phylogenetic tree was further calculated between the three aging strategies and Contr treatment. The results showed that the manipulation of straw and the addition of a decomposing microbial consortium significantly increased the abundance of a range of species (42 species) belonging to Firmicutes during hydrochar aging. These species mentioned above were mainly from guild 6 (79%), and their genus-level classifications were mainly *Bacillus* (17 species) and *Ornithinibacillus* (3 species). It has been widely suggested that members of the genus *Bacillus,* as core microbial taxa in most composts, perform multiple functional activities, including macromolecular substrate degradation (Zhang et al., 2023) and P fraction mobilization (Zhang et al., 2021a). In addition, cyclic lipopeptides from *Bacillus* spp. have been introduced to prevent abiotic and biotic stresses and promote plant growth (Tunsagool et al., 2023). The evidence suggested that *Ornithinibacillus* is associated with ammonium nitrogen and plays a regulatory role in the nitrification process during composting (Zhai et al., 2023). It can be concluded that *Bacillus* and their related species play an important role in substrate degradation and nutrient cycling during hydrochar aging.

Furthermore, the results of the functional prediction analysis demonstrated that these species enriched in the SJ treatment harboured higher potential in nitrogen assimilation and organic phosphorus mineralization/transport-associated functions than other species in guilds (Supplementary Data 1). Most microorganisms rely on biologically available forms of nitrogen for growth and metabolism (Liu et al., 2023; Morrissey et al., 2018). As anticipated from previous work, N assimilation is practised by many microbial taxa (Morrissey et al., 2018). Therefore, the community composition of N assimilation microorganisms could depend primarily on specific environmental conditions rather than phylogeny. Food waste is relatively rich in organophosphorus compounds (Kumar et al., 2023). The main processes of P transformations during composting are the absorption of P by the phosphate inorganic transporter (Pit) and specific transporter (Pst) systems, the dissolution of inorganic P, and the mineralization of organo-phosphorus compounds (Bergkemper et al., 2016). In addition, microbial populations related to cellular P transport can compete with other organisms in composting for available P because they have efficient phosphate uptake systems (Bergkemper et al., 2016). This finding indicated that microbes with P transformation-associated functions could be prevalent during hydrochar aging processes. This is, however, still to be verified, as the functional output from PICRUSt2 analysis is less likely to resolve rare environment-specific functions (Douglas et al., 2020). In addition, this study was conducted under laboratory conditions, and the effectiveness of straw and microbial inoculum addition in the aged hydrochar production should be further determined. In a future study, it is also necessary to investigate the promoting effects of aged hydrochar on plant growth, as well as their long-term impacts on soil and environmental health.

## 4. Conclusion

This work revealed that the addition of straw and efficient-degrading microbes can increase the reaction temperature by ∼13%, and accelerate the process by ∼30% of hydrochar aging. Bacterial communities, rather than fungal communities, dominated throughout aging, as evidenced by their dramatically higher diversity and higher proportion in the microbial interaction network. The dynamics of nutrient contents during hydrochar aging process, especially nitrogen and phosphorus, can be attributed to the stimulation of the key bacterial guilds, mainly including the *Bacillus-*like species. In a future study, the promoting effects of aged hydrochar applications on plant growth, as well as its long-term impacts on soil and environmental health are need to be investigated. This study demonstrates the feasibility of artificially regulating microbial communities to accelerate the aging processes, contributing to the development in agricultural applications of hydrochar.

## CRediT authorship contribution statement

**Yang Ruan:** Writing – original draft, Writing – review & editing, Visualization, Formal analysis. **Ziyan Wang:** Experimental investigation. **Shiyong Tan**: Validation, Funding acquisition. **Hao Xu:** Experimental investigation, Writing – review & editing. **Liyue Wang:** Formal analysis. **Lixuan Ren:** Supervision. **Zhipeng Liu:** Methodology, Writing – review & editing. **Shiwei Guo:** Data curation, Writing – review & editing. **Qirong Shen:** Conceptualization. **Guohua Xu:** Resources, Conceptualization, Funding acquisition. **Ning Ling:** Writing – review & editing, Supervision.

## Data availability

The raw sequence data reported in this paper have been deposited in the Genome Sequence Archive (Genomics, Proteomics & Bioinformatics 2021) in National Genomics Data Center (Nucleic Acids Res 2022), China National Center for Bioinformation / Beijing Institute of Genomics, Chinese Academy of Sciences (GSA: CRA012057) that are publicly accessible at https://ngdc.cncb.ac.cn/gsa.

## Acknowledgments

This work was supported by the National Nature Science Foundation of China (41961124005); and the Science and Technology Innovation Program of Hunan Province (2022RC3057). We thank Kang Chen and Yide Shan for the experimental investigation, sampling work and sample processing assistance.

