## supplementary materials for "Microbial community assembly during aging of food waste-derived hydrochar: key bacterial guilds mediate nutrient dynamics"

a, Jiangsu provincial key lab for solid organic waste utilization, Key lab of organic-based fertilizers of China, Jiangsu Collaborative Innovation Center for Solid Organic Wastes, Educational Ministry Engineering Center of Resource-saving fertilizers, Nanjing Agricultural University, Nanjing 210095, China.

b, Centre for Grassland Microbiome, State Key Laboratory of Grassland Agro-ecosystems, College of Pastoral Agricultural Science and Technology, Lanzhou University, Lanzhou, 730020, Gansu, China.

c, Hunan Biological Fertilizer Engineering Technology Research Center, Changsha 410083, China.

*Corresponding author:

Ning Ling

Zhipeng Liu

**Supplementary Information includes:**

Table S1

Figures S1-S2

**Table S1 The alpha diversity of bacterial and fungus communities under four aging treatments during the aging process.**

| Treatment | Aging time | Richness | | Shannon index | |
|---|---|---|---|---|---|
| | | Bacteria | Fungi | Bacteria | Fungi |
| Contr | Day 1 | 2225±46 a | 146±47 a | 7.2±0.5 a | 2.4±0.3 a |
| | Day 7 | 2400±316 a | 76±15 a | 5.5±1.8 a | 1.6±0.7 ab |
| | Day 14 | 1638±120 a | 105±81 a | 5±0.1 a | 2±0.6 ab |
| | Day 35 | 3246±1315 a | 100±8 a | 8.4±2.1 a | 0.9±0.2 b |
| S | Day 1 | 3504±1161 a | 77±10 a | 8.8±2.1 a | 0.6±0.1 b |
| | Day 7 | 2182±347 a | 74±16 a | 6.4±0.4 ab | 1.8±0.8 ab |
| | Day 14 | 2337±155 a | 98±15 a | 6.5±0.9 ab | 2.7±0.3 a |
| | Day 35 | 1754±174 a | 81±6 a | 4.3±1.7 b | 1.4±0.3 b |
| ST | Day 1 | 2808±1014 a | 85±27 a | 7.7±2.2 a | 1.3±0.8 a |
| | Day 7 | 2804±962 a | 76±7 a | 5.8±3.5 a | 2.1±0.1 a |
| | Day 14 | 2052±786 a | 96±20 a | 6.2±2.2 a | 0.8±0.4 a |
| | Day 35 | 1781±47 a | 144±64 a | 5.7±0.1 a | 1±0.1 a |
| SJ | Day 1 | 3034±1066 a | 79±14 a | 7.6±2.2 a | 1.2±0.6 ab |
| | Day 7 | 2212±610 ab | 85±13 a | 7±0.4 a | 1.4±0.7 b |
| | Day 14 | 1967±46 ab | 87±20 a | 6±0.2 a | 1.7±0.5 a |
| | Day 35 | 1565±144 b | 61±35 a | 5.3±1.3 a | 1.7±1.2 ab |

Note: Contr represents the normal aging treatment; S represents the hydrochar aging with straw addition; ST represents the hydrochar aging with straw and decomposition agent addition; SJ represents the hydrochar aging with straw and the microbial consortium addition. Letters: significant differences among aging periods at the same aging treatment.

### ***Supplementary Figures:***

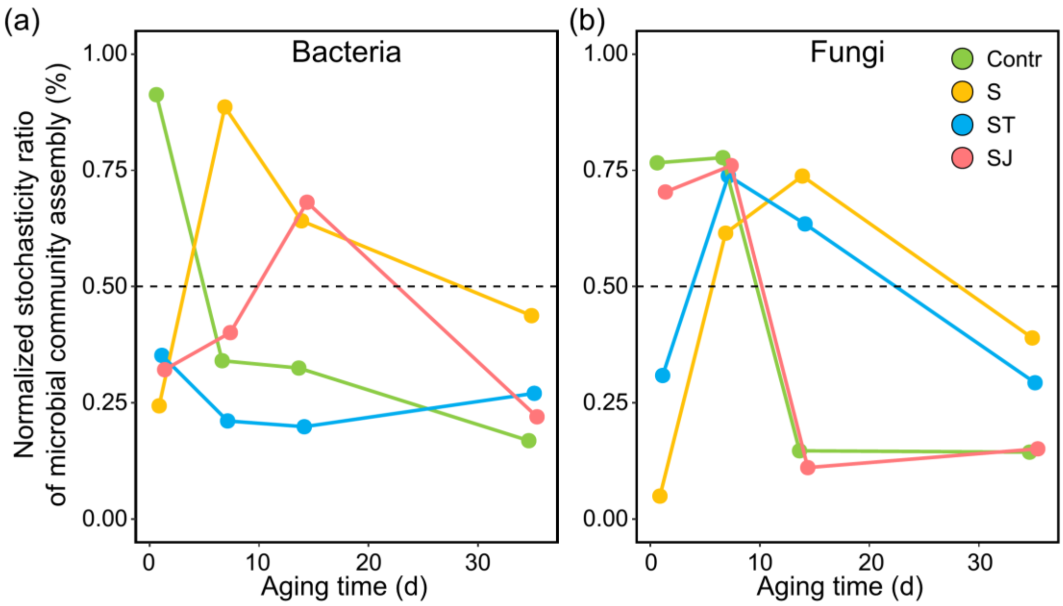


**Fig. S1 Dynamic changes in the normalized stochasticity ratio (NST) during the succession of the hydrochar bacterial (a) and fungal (b) communities under four aging strategies.** NSTs greater than 50% are considered to be dominated by stochastic processes; NSTs less than 50% are considered to be dominated by deterministic processes. Colours represent different aging strategies. Contr represents the normal aging treatment; S represents the hydrochar aging with straw addition; ST represents the hydrochar aging with straw and decomposition agent addition; SJ represents the hydrochar aging with straw and the microbial consortium addition. Letters: significant differences among aging periods at the same aging treatment.

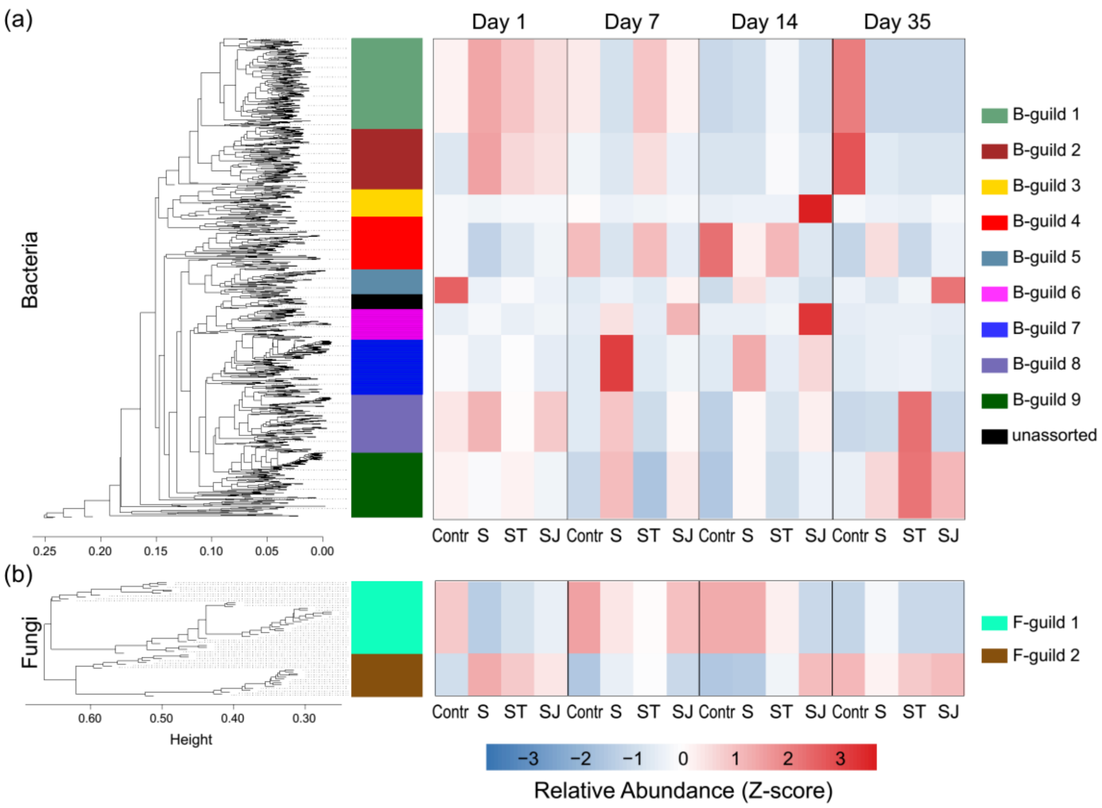


**Fig. S2 Determination of bacterial (a) and fungal (b) guilds using WGCNA.** This analysis was based on the fold change of the differential abundance species ($Log_2FC$) in the comparison between each aging strategy and the normal aging treatment during the aging periods. The heatmap shows the average relative abundance of guilds (Z score normalized). Contr represents the normal aging treatment; S represents the hydrochar aging with straw addition; ST represents the hydrochar aging with straw and decomposition agent addition; SJ represents the hydrochar aging with straw and the microbial consortium addition. Letters: significant differences among aging periods at the same aging treatment.
